# Longitudinal Trajectories of Cognition and Neural Metrics as Predictors of Persistent Distressing Psychotic-Like Experiences Across Middle Childhood and Early Adolescence

**DOI:** 10.1101/2025.01.20.633817

**Authors:** Nicole R. Karcher, Fanghong Dong, Emma C. Johnson, Sarah E. Paul, Can Misel Kilciksiz, Hans Oh, Jason Schiffman, Arpana Agrawal, Ryan Bogdan, Joshua J. Jackson, Deanna M. Barch

## Abstract

**Objectives:** Psychotic-like experiences (PLEs) may arise from genetic and environmental risk leading to worsening cognitive and neural metrics over time, which in turn lead to worsening PLEs. Persistence and distress are factors that distinguish more clinically significant PLEs. Analyses used three waves of unique longitudinal Adolescent Brain Cognitive Development Study data (ages 9-13) to test whether changes in cognition and structural neural metrics attenuate associations between genetic and environmental risk with persistent distressing PLEs.

**Methods:** Multigroup univariate latent growth models examined three waves of cognitive metrics and global structural neural metrics separately for three PLE groups: persistent distressing PLEs (n=356), transient distressing PLEs (n=408), and low-level PLEs (n=7901). Models then examined whether changes in cognitive and structural neural metrics over time attenuated associations between genetic liability (i.e., schizophrenia polygenic risk scores/family history) or environmental risk scores (e.g., poverty) and PLE groups.

**Results:** Persistent distressing PLEs showed greater decreases (i.e., more negative slopes) of cognition and neural metrics over time compared to those in low-level PLE groups. Associations between environmental risk and persistent distressing PLEs were attenuated when accounting for lowered scores over time on cognitive (e.g., picture vocabulary) and to a lesser extent neural (e.g., cortical thickness, volume) metrics.

**Conclusions:** Analyses provide novel evidence for extant theories that worsening cognition and global structural metrics may partially account for associations between environmental risk with persistent distressing PLEs.

---

Psychotic-like experiences (PLEs) are subclinical abnormal thought content and perception abnormalities which are on the continuum of psychosis spectrum(1,2). Studies indicate evidence for the clinical relevance of distressing PLEs, with 57-92% of those with *significantly* distressing PLEs developing a diagnosable mental health problem later in adulthood(3,4), including psychotic disorders or other diagnosable conditions, such mood and behavioral disorders(3,5). Although a recent literature review indicated some research finds PLEs are not associated with sustained negative outcomes(2), other research finds childhood and adolescent PLEs are associated with a range of impairments in domains including cognition and pathophysiology, including structural neural metrics(6–10). Our recent work using the Adolescent Brain Cognitive Development^℠^ (ABCD) Study data found evidence that persistent (i.e., occurring over multiple years) and distressing PLEs are associated with a range of risk factors (e.g., environmental exposures, cognitive impairments) measured at the start of adolescence(10). This present work represents an important expansion from prior cross-sectional work by examining the extent to which these identified cognitive and structural neural factors over time represent mechanisms underlying associations between genetic and environmental risk with persistent distressing PLEs.

Literature shows that longitudinal changes in cognition, including executive function, are both risk factors for and consequences of general psychopathology(11). There is evidence that longitudinal changes in cognition may be particularly associated with the development of schizophrenia spectrum symptoms(12), and that decline in cognition is present at the subclinical phase of psychosis spectrum in early ages(13). Although results on cognitive deficits among high risk or prodromal populations are promising for early detection of psychosis, there is limited data about changes in cognitive domains in populations experiencing PLE even before the risk phase(12), with some evidence for development lags in cognition(13), although other evidence for weak declines over time(14). Following changes in cognition may be used as an important tool for tracking worsening functioning.

Early childhood through adolescence is a key developmental period for brain development and maturation. Brain changes during this time include but are not limited to general decreases in cortical thickness, surface area, and volume(15). This peak of brain maturation and neuroplasticity during adolescence makes this a period of vulnerability for the development of psychopathology(16). Studies examining longitudinal changes in structural neural metrics associated with the psychosis symptom spectrum find individuals who convert to psychosis generally show reductions in grey matter in a range of frontal(17,18)), temporal(17,19)), and cingulate and cerebellar regions(19)). One recent review indicates that this accelerated decline in a range of areas (e.g., frontal, temporal, cingulate, and parietal) extends to individuals experiencing PLEs(20)). Overall, this research indicates psychosis spectrum symptoms are associated with longitudinal structural neural deviations spanning a diverse range of neural regions.

Theory indicates that PLEs may arise as a result of a combination of genetic and environmental factors leading to worsening pathophysiological factors (e.g., cognition, structural neural metrics), which in turn lead to worsening PLEs(21)). The present work therefore also explored whether cognitive or neural metrics over time attenuated associations between genetic (quantified using schizophrenia polygenic scores and family history of psychosis) and environmental risk to persistent distressing PLEs. Our previous work found evidence that significantly distressing PLEs are associated with genetic liability (as measured by polygenic risk scores for schizophrenia, PLEs, broad cross-disorder psychopathology, and lowered educational attainment)(22), as well as environmental risk (as measured by a composite of previously implicated environmental risk factors (i.e., poverty, lack of neighborhood safety, deprivation, and estimated lead risk exposure)(10,23). Showing changes in cognitive and neural metrics may alter links between genetic and environmental factors with worsening PLEs informs both our understanding of the pathogenesis of PLEs and the possibility that altered cognitive and neural metrics may affect PLEs trajectories, information useful for early identification and intervention (e.g., cognitive therapy and remediation(24)) efforts.

The Adolescent Brain Cognitive Development (ABCD) study is a longitudinal study that aims to understand the neurodevelopmental trajectories of a diverse sample of over 11,000 children across the United States. The present work provides several important novel advances towards understanding the pathogenesis of early PLEs including: a) providing novel evidence for longitudinal changes in cognitive and structural neural metrics predicting persistent distressing PLEs, something that has not previously been examined in any dataset, including ABCD data, b) providing novel evidence for whether associations between genetic and environmental factors and PLEs are attenuated by cognition and neural metrics over time, consistent with a plausible mediation pathway. It was hypothesized that impairments in cognitive and deviations of global structural metrics over time would be most strongly associated with persistent distressing PLEs compared to low-level PLEs or transient distressing PLEs, and that these metrics would attenuate associations between both genetic and environmental factors with PLEs.

## Methods

### Participants

The ABCD Study is a large-scale study tracking 9-10-years-olds recruited from 21 research sites across the United States (Supplement for study-wide exclusion details). ABCD Data Release 5.0 (DOI:<u>10.15154/z563-zd24</u>) includes 5 waves of data: baseline (N=11,868), 1-year follow-up (N=11,217), 2-year follow-up (N=10,973), 3-year follow-up (N=10,331), and a portion of the 4-year follow-up (N=4,754; Table 1 for sample characteristics). Each participant’s past 3 waves of data were used to define PLE groups (see below). Cognitive and MRI data was obtained at baseline, 2-year follow-up and 4-year follow-up. All available data were used in analyses and missing data were handled using full information maximum likelihood (fiml) estimation (see Supplement for models using imputed data; Supplemental Table 1 for sample sizes for included variables).

**Table 1.**
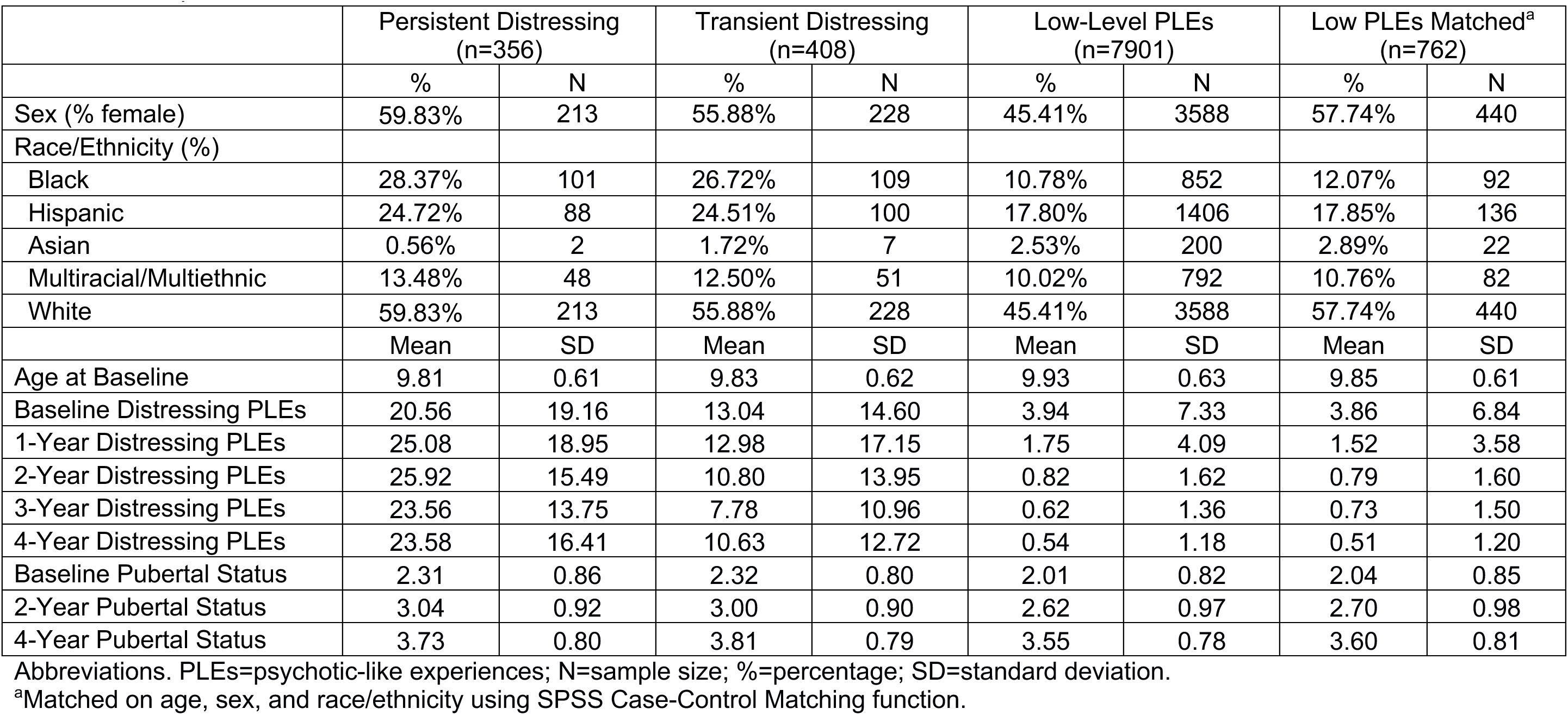
Participant Characteristics.

### Measures

#### Prodromal Questionnaire-Brief Child Version (PQ-BC)

Participants completed the previously validated Prodromal Questionnaire-Brief Child Version (PQ-BC)(9)). As the present work followed up our prior cross-sectional work, we used the same PLE group designations as in our prior work(10,25), with group definitions based on previous research(26). Consistent with this previous research, distress scores were calculated as the total number of 21 questions endorsed weighted by level of distress [i.e., 0=no, 1=yes (but no distress), 2-6=yes (1+score on distress scale); range:0-126]. Analyses examined the following groups (Table 1): 1) a *persistent distressing* PLEs group that scored>=1.96SDs above the mean for distressing PLEs for 2 or more of the 3 waves of data based on the participant’s last three waves of data from baseline through 4-year follow-up (n=356, range of mean PLEs across waves: 20.56-25.92(27); means and standard deviation were examined across the entire sample for each assessment wave); 2) a *transient distressing* PLEs group that scored >=1.96 SDs above the mean for distressing PLEs for 1 wave of data and scored <=0.50 SDs above the mean for distressing PLEs for the other 2 waves of data (n=408, range of mean PLEs across waves: 7.78-13.04); 3) a *low-level PLEs* group that scored <=0.50SDs above the mean for distressing PLEs for all three waves of data (n=7901, range of mean PLEs across waves: 0.54-3.94).

#### Neuropsychological Test Battery

National Institutes of Health Toolbox Cognitive Battery (NIHTB-CB) tests included flanker inhibitory control, pattern processing speed, picture vocabulary, and readings tests, administered at baseline, 2-year-follow-up and 4-year follow-up assessments, as well as list sorting working memory, which was administered at baseline and 4-year follow-up (for included tests, administration time is approximately 22 minutes; for more details, see(27).

#### Structural MRI Measures

Consistent with prior work(10), structural MRI measures included intracranial volume, total cortical and subcortical volumes(28), total surface area(29), and mean cortical thickness(30). All data were acquired on a 3T scanner (Siemens, General Electric, or Phillips) with a 32-channel head coil and completed T1-weighted and T2-weighted structural scans (1mm isotropic). Structural neuroimaging processing was completed using FreeSurfer version 5.3.0 through standardized processing pipelines(31) (Supplement for additional details).

Participants that did not pass FreeSurfer Quality Control measures (i.e., at least one T1 scan that passed all quality control metrics) were excluded from analyses (n=142). We used longitudinal ComBat harmonization(32), with age and sex added as biological covariates to the design matrix, and individual, site, and family nested as a random effects, to estimate and remove individual scanner effects (i.e., using Siemens, Phillips, GE device serial numbers) from MRI measures prior to entry into models.

#### Genetic and Environmental Risk Scores

To account for genetic liability, we examined schizophrenia polygenic scores (PGS)(22) and family history of psychosis. Summary statistics from well-powered discovery GWASs of schizophrenia for youth most genetically similar to individuals from European reference populations (N=69,369 cases+236,642 controls)(33) and youth most genetically similar to individuals from African reference populations (N=6152 cases+3918 controls)(34) were used. PGS were generated using polygenic risk scores-continuous shrinkage (PGS-CS; see Supplement for more information)(35). These analyses were done in a subset of individuals from European ancestry (n=4533) and from African ancestry (n=1201), due to the sample compositions of the discovery GWASs and evidence that polygenic risk scores do not translate well across genetic ancestries in an unbiased manner(36). To examine a proxy for potential genetic (and non-genetic) liability across the entire ABCD sample, we also examined whether caregivers endorsed any family history of psychosis. For environmental risk scores, based on previous work(10,23) we created a principal component analysis (PCA) using baseline metrics of caregiver-rated perception of neighborhood safety, number of years at current residence, and based on primary address: drug crime exposure, overall deprivation, rate of poverty, and lead exposure risk estimates (see Supplement). The extracted first principal component was used as a predictor in analyses.

#### Statistical Analysis

Multi-group (i.e., persistent distressing, transient distressing, low-level PLEs) growth curve models (growth function in lavaan package in R version 0.6-15) estimated slopes for cognitive and global structural neural metrics over time (i.e., baseline, 2-year-follow-up and 4-year follow-up), simultaneously modeling and freely estimated for each of the three PLE groups. Differences between the groups were specified in each model as difference scores between individual parameter estimates. Given the focus of the present work on understanding persistent distressing PLEs, analyses primarily focused on the slope difference of persistent distressing vs. low-level PLEs. To examine specificity of findings, analyses additionally examined transient distressing vs. low-level PLEs and persistent distressing vs. transient distressing PLEs. To account for multiple comparisons, these difference between slope estimates were false discovery-rate (FDR) corrected (11 FDR-corrected tests: 6 cognition metrics, 5 MRI metrics) for each group comparison. Models also examined mean initial scores (i.e., as indexed by the intercept which was set at the baseline assessment) separately for each of the three PLE groups. Analyses included sex at baseline as a time invariant covariate, with pubertal status and age in months as time varying covariates. Variables were examined for skew, and all continuous variables with skew ≥|1.96| were winsorized to 3SDs.

Follow-up analyses examined whether changes in cognition and neural metrics over time attenuate associations between genetic and environmental risk with persistent distressing PLEs, a result that would be consistent with the possibility that changes in cognition or neural metrics are mediators of this association. First, we examined associations between genetic liability (PGS, family history of psychosis) and, separately, environmental risk and PLE group comparisons. For significant genetic or environmental factors, we examined whether their association with PLE group comparisons were attenuated when including cognition and structural neural metric slopes using the lavaan package growth function in R (these models included the same covariates as in original analyses). Analyses were FDR corrected across 11 tests.

## Results

Information about the persistent distressing, transient distressing, and low-level PLE groups can be found in Table 1.

### Cognition Trajectories in PLE groups

As can be seen in Figure 1, the persistent distressing PLEs group relative to the low-level PLEs group showed lower initial scores (i.e., negative intercepts; Supplemental Table 2) and worsening performance over time (i.e., negative slopes) across cognitive tasks (difference between slope estimates≤-0.093, FDR*p*s<.02, except for Flanker inhibitory control: difference between slope estimates=-0.141, FDR*p*=.07; see Table 2 for group differences; see Supplemental Table 3 for estimates for each PLE group). The transient distressing PLEs group showed lower initial values across tasks (i.e., negative intercepts; Supplemental Table 2) and significant worsening performance over time (i.e., negative slopes; Figure 1) for the picture vocabulary, picture sequence memory, and working memory tasks compared to the low-level PLEs groups (difference between slope estimates≥-0.16, FDR*p*s<.01).

**Figure 1.**
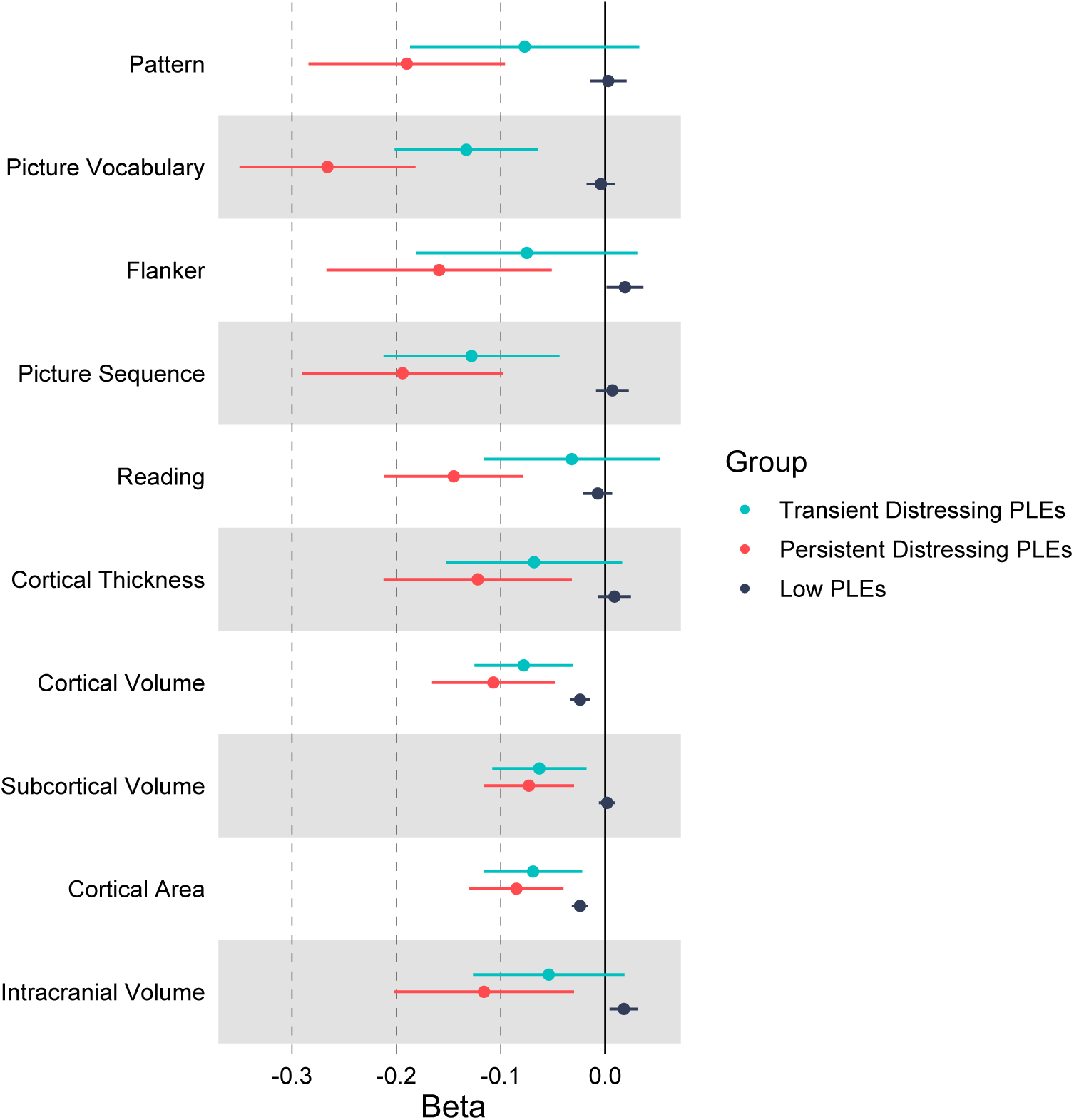
Cognitive and neural metric slope estimates (z-scored across the entire sample) for low-level psychotic-like experiences (PLEs), persistent distressing PLEs, and transient distressing PLEs groups when including covariates (i.e., age, sex, pubertal status).

**Table 2.** Difference between slope estimates Comparisons of Each of the PLE Groups.

|  | Persistent Distressing vs. Low-Level PLEs |  |  |  |  | Transient Distressing vs. Low-Level PLEs |  |  |  |  | Persistent Distressing vs. Transient Distressing PLEs |  |  |  |  |
| --- | --- | --- | --- | --- | --- | --- | --- | --- | --- | --- | --- | --- | --- | --- | --- |
|  | Est. | lower 95% CI | upper 95% CI | Z value | FDRp | Est. | lower 95% CI | upper 95% CI | Z value | FDRp | Est. | lower 95% CI | upper 95% CI | Z value | FDRp |
| Pattern | -0.181 | -0.287 | -0.075 | -3.342 | 0.002 | -0.083 | -0.196 | 0.031 | -1.433 | 0.17 | 0.098 | -0.056 | 0.252 | 1.251 | 0.88 |
| Picture Vocabulary | -0.171 | -0.256 | -0.086 | -3.934 | <.001 | -0.168 | -0.229 | -0.107 | -5.417 | <.001 | 0.003 | -0.1 | 0.106 | 0.059 | 0.95 |
| Flanker | -0.141 | -0.293 | 0.011 | -1.82 | 0.07 | -0.115 | -0.24 | 0.011 | -1.791 | 0.09 | 0.026 | -0.169 | 0.222 | 0.265 | 0.94 |
| Picture Sequence | -0.173 | -0.283 | -0.063 | -3.08 | 0.003 | -0.16 | -0.24 | -0.08 | -3.908 | <.001 | 0.013 | -0.122 | 0.147 | 0.187 | 0.94 |
| Reading | -0.093 | -0.168 | -0.018 | -2.442 | 0.02 | -0.075 | -0.149 | 0 | -1.969 | 0.07 | 0.018 | -0.085 | 0.122 | 0.35 | 0.94 |
| List Sorting | -0.28 | -0.484 | -0.077 | -2.698 | 0.009 | -0.198 | -0.348 | -0.049 | -2.602 | 0.01 | 0.082 | -0.167 | 0.331 | 0.645 | 0.94 |
| Thickness | -0.144 | -0.055 | -0.233 | -3.169 | 0.003 | -0.046 | -0.032 | -0.124 | -1.156 | 0.25 | 0.098 | -0.018 | 0.213 | 1.656 | 0.88 |
| Volume | -0.175 | -0.107 | -0.244 | -5.026 | <.001 | -0.122 | -0.062 | -0.182 | -3.992 | <.001 | 0.054 | -0.036 | 0.143 | 1.177 | 0.88 |
| Subcortical Volume | -0.11 | -0.051 | -0.169 | -3.67 | <.001 | -0.1 | -0.05 | -0.149 | -3.949 | <.001 | 0.011 | -0.065 | 0.086 | 0.278 | 0.94 |
| Surface Area | -0.115 | -0.048 | -0.182 | -3.349 | 0.002 | -0.096 | -0.045 | -0.147 | -3.689 | <.001 | 0.019 | -0.064 | 0.102 | 0.444 | 0.94 |
| Intracranial Volume | -0.162 | -0.045 | -0.28 | -2.702 | 0.009 | -0.11 | -0.028 | -0.192 | -2.623 | 0.01 | 0.052 | -0.089 | 0.194 | 0.726 | 0.94 |

Abbreviations. Est=estimate; PLE=psychotic-like experiences; 95% CI= 95% confidence interval; Z=Z statistics; $p$ =p-value; FDR $p$ =false discovery-rate p-value

### MRI Metrics Trajectories in PLE groups

For structural neural metrics, as can be seen in Figure 1, the persistent distressing group relative to the low-level PLEs group showed lower initial values (i.e., negative intercepts; Supplemental Table 2) and accelerated decreases over time (i.e., negative slopes) across all metrics, including cortical and subcortical volume, cortical surface area, cortical thickness, and intracranial volume (difference between slope estimates≤-0.11, FDR*p*=.009, Table 2; see Supplemental Table 3 for estimates for each PLE group). The transient distressing PLEs group relative to the low-level PLEs group showed lower initial values (i.e., negative intercepts; Supplemental Table 2) and accelerated decreases over time (i.e., negative slopes, Figure 1) for cortical and subcortical volume, cortical area, and intracranial volume, but not for cortical thickness.

### Follow-up Analyses

See Supplement for results using latent profile analyses, with results highly consistent. See Supplemental Table 4 for models in a low-level PLEs sample matched to the PLEs groups in age, sex, and race/ethnicity using SPSS Case/Control matching procedure, with results remaining consistent. As seen in Supplemental Table 5, results remain consistent when including youth-reported internalizing and externalizing symptoms over time.

### Models Testing Attenuation of Associations between Genetic and Environmental Factors and PLE Groups

Greater environmental risk scores and greater family history of psychosis (FDRps<.05; Supplemental Table 6), but not schizophrenia PGS scores, were associated with being in the persistent distressing PLE versus the low-level PLEs group. When family history and environmental risk scores were entered simultaneously, only environmental risk scores remained associated with the persistent distressing PLEs (95%CI:0.174,0.311).

### Genetic and Environment Risk with Cognition and Imaging Slope Metrics

Given the associations between both family history of psychosis and environmental risk with PLE groups, we examined associations between environmental risk scores and family history with cognition and imaging slope metrics. Greater environmental risk scores were associated with lower cognitive scores over time (i.e., negative slopes; βs<-.08, FDRps<.004; Supplemental Table 7, except for pattern processing speed).

Higher environmental risk scores were associated with lower imaging slopes (βs<-0.08, FDRps<.001) across all imaging metrics (i.e., thickness, cortical volume, subcortical volume, cortical area, and intracranial volume; Supplemental Table 7). Family history of psychosis was not strongly associated with any of the cognitive or imaging scores over time (FDRps>.10).

### Evidence for Cognitive and Neural Trajectories Attenuating Environmental Risk with PLEs

Given the above results, to explore whether slopes of cognition and imaging metrics attenuated associations between environmental risk with PLEs, associations between each of the significantly associated environmental risk and cognition/imaging slope measures with PLE groups were examined. As can be seen in Figure 2, the inclusion of cognitive slopes and to a lesser extent imaging slopes attenuated associations between environmental risk with persistent distressing PLEs (i.e., slopes attenuated between 2.4% (intracranial volume) – 26.38% (picture vocabulary)).

**Figure 2.**
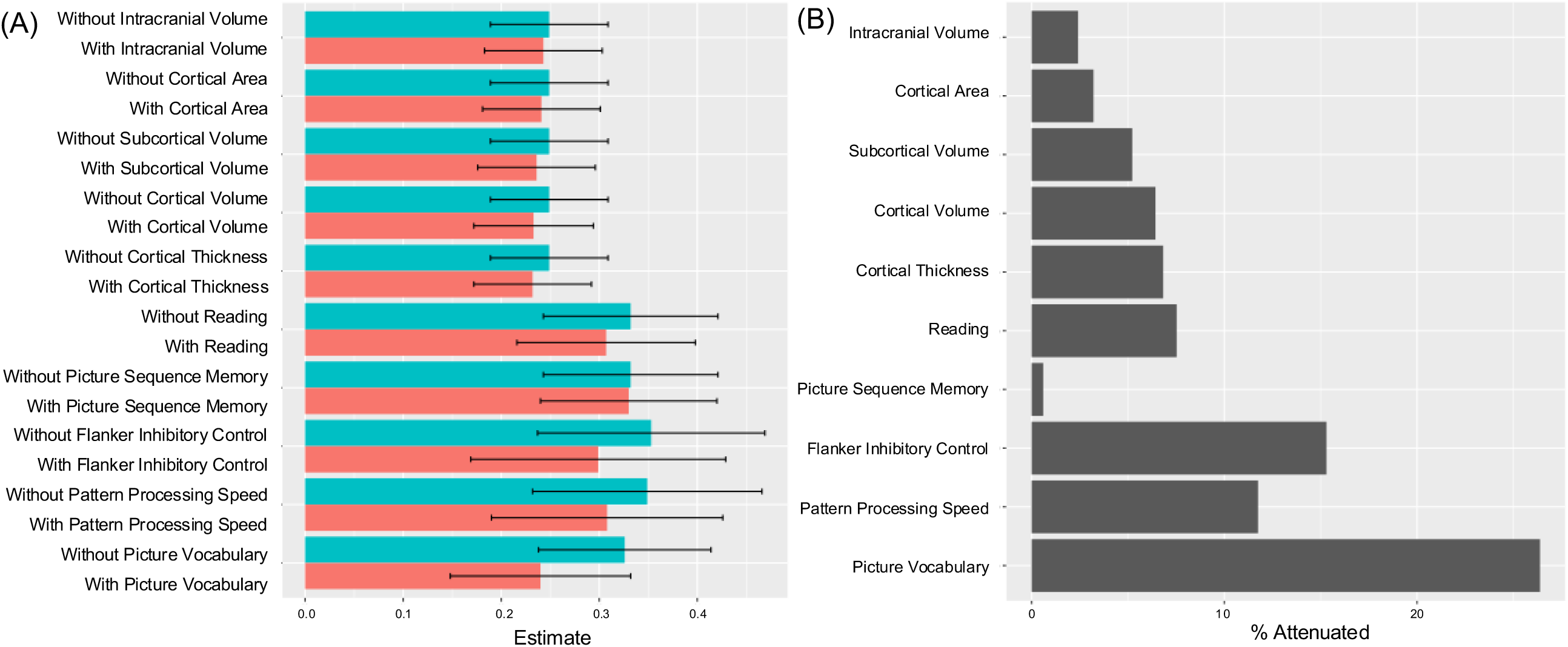
(A) Estimates for models examining associations between environmental risk with Persistent Distressing PLEs vs. Low-level PLEs either with or without accounting for cognitive or imaging metric slopes. (B) Estimated proportion of the association between environmental risk with Persistent Distressing PLEs vs. Low-level PLEs attenuated when accounting for each cognitive or imaging metric slope metric.

## Discussion

The present study examines adolescent trajectories of cognition and structural neural metrics over the course of approximately six years in relation to persistent and transient distressing PLEs. Deviations in cognitive and cortical thickness metrics were evident in a gradient across PLE groups, with persistent distressing PLEs compared to low-level PLEs showing the largest impairments, followed by the transient distressing PLEs group, even when accounting for youth-reported internalizing and externalizing symptoms (Supplemental Table 5). Although the effect sizes were smaller than those generally seen in clinical high-risk studies(12), they were larger than previous cross-sectional work in this sample(10), providing evidence for the importance of understanding declines over time. Further, worsening cognitive and to a lesser degree imaging metrics attenuated links between higher environmental risk and persistent distressing PLEs. The findings provide key information that youth who show persistent distressing PLEs are characterized by demonstrable decreases in cognition and neural metrics, and that these impairments in cognition and structural metrics may partially account for associations between environmental risk with persistent distressing PLEs.

Across cognitive tests, the distressing PLEs groups showed worsening performance over time, with the strongest findings for persistent distressing PLEs. The persistent distressing PLEs group showed evidence of worsening performance over time across cognitive metrics, whereas the transient distressing PLEs group showed worsening performance in a subset of tests including picture vocabulary, picture sequence, and working memory. These results are generally consistent with previous cross-sectional work, including our work in the ABCD study(10), where findings have pointed to broad ranging cognitive impairments associated with PLEs(10,37) spanning both crystallized and fluid cognition impairments, with effect strongest for persistent distressing PLEs.

For global structural imaging metrics, as with the cognitive metrics, the distressing PLEs groups showed broad accelerated declines over time, with the strongest findings for persistent distressing PLEs. Although volume, area, and thickness generally decline across adolescence(15) especially the persistent distressing PLEs group showed evidence of accelerated declines in cortical metrics over time compared to the low-level PLEs group. Persistent distressing PLEs showing evidence of greater reductions in cortical thickness over time is consistent with prior work indicating that accelerated cortical thinning often precedes the onset of psychosis(17). Overall, although the persistent distressing PLEs group showed widespread accelerated declines in global structural metrics over time, there was some evidence for specificity, including that only the persistent distressing PLEs group showed accelerated declines in cortical thickness when compared to the low-level PLEs group.

Attenuation analyses point to the potential role of worsening cognition and brain metrics in linking environmental risk to persistent distressing PLEs. Genetic liability analyses indicated that family history of psychosis was associated with persistent distressing PLEs, but polygenic scores for schizophrenia were not. Transient distressing PLEs were not strongly associated with family history of psychosis or schizophrenia polygenic scores, which may point the heterogeneity of transient distressing PLEs. Consistent with our prior work, persistent distressing PLEs were strongly associated with greater environmental risk scores(23), bolstering evidence for the importance of environmental factors including poverty and adverse life events in relation to persistent PLEs(23,38). Worsening cognitive and neural metrics over time attenuated associations between environmental risk and persistent distressing PLEs, indicating that the robust link of the environment with PLEs may be through diffuse worsening of cognitive and neural metrics over time(42). Overall, these data provide important empirical evidence consistent with previous theoretical models regarding the development and worsening of PLEs(39), namely that both genetic liability and environmental risk play a role in adolescents experiencing persistent distressing PLEs, and these environmental associations are attenuated by worsening cognition and by worsening neural metrics.

This work had several limitations and points to consider. The groups were created based on *a priori* (versus data driven) definitions of group membership, as we were specifically interested in following up on our previous work examining these definitions of PLEs(10), although it is likely that heterogeneity exists within these groups (e.g., remitting trajectories), as evidenced by the greater variability in cognitive and neural metrics slopes for PLE groups compared to the low-level PLE group (see Figure 1). Further, latent profile analyses (Supplemental Table 8) generally supported these results, with 98% of the persistent distressing group falling within the high PLEs latent class, whereas only 31% of the transient distressing PLEs group fell into the high PLEs latent class. Research should examine the influence of factors that may vary by marginalized communities, including structural racism, as well as service utilization and potential targets for intervention, including physical health-related variables (e.g., sleep, physical activity, nutrition, substance use) in interrelationships between environment, cognitive, neural metrics, and PLEs over time. Variables chosen for analyses relied on our past cross-sectional work examining genetic liability, cognition, structural metrics, and environment(10,22,23). Polygenic scores models were limited to ancestries to ancestral populations where we have previously validated PGS. Important genetic effects on PLEs in individuals of other genetic ancestries might have been missed, however, this is likely partially recovered through our family history measure which in part assesses genetic liability. Genetic liability analyses therefore have incomplete generalizability, which may have the potential to increase structural inequities including health disparities (40), necessitating future research to understand associations between genetic liability, structural inequities, and risk for psychosis. Future work should examine the role of other potentially influential factors, including drug use and mental health service utilization in potentially exacerbating or attenuating these associations, respectively.

Overall, there was evidence that compared to the low-level PLEs group, persistent distressing PLEs showed broad evidence of worsening of cognitive functioning and accelerated declines in global structural neural metrics over the course of approximately six years. The present work provides important evidence for extant theory, providing some of the first evidence for general cognitive and global structural neural metrics attenuate links between environment with worsening psychosis spectrum symptoms over the course of middle childhood and adolescence. This research provides important advances over previous cross-sectional work (Supplemental Table 9), indicating that declines in cognitive and neural metrics are already evident in middle childhood, declines that are generally smaller than in high-risk populations(12), and therefore this developmental time period may represent a key period for early intervention. This work provides information regarding the nature of worsening PLEs in adolescence, including the role of cognition and the brain as potential mechanisms linking genes and environment to worsening PLEs.

## Supporting information

Supplemental Methods and Results

## Acknowledgments

Data used in the preparation of this article were obtained from the Adolescent Brain Cognitive Development (ABCD) Study (https://abcdstudy.org), held in the NIMH Data Archive (NDA). This is a multisite, longitudinal study designed to recruit more than 10,000 children age 9–10 and follow them over 10 years into early adulthood. The ABCD Study is supported by the National Institutes of Health and additional federal partners under award numbers U01DA041022, U01DA041025, U01DA041028, U01DA041048, U01DA041089, U01DA041093, U01DA041106, U01DA041117, U01DA041120, U01DA041134, U01DA041148, U01DA041156, U01DA041174, U24DA041123, and U24DA041147. A full list of supporters is available at https://abcdstudy.org/nih-collaborators. A listing of participating sites and a complete listing of the study investigators can be found at https://abcdstudy.org/principal-investigators.html. ABCD consortium investigators designed and implemented the study and/or provided data but did not necessarily participate in analysis or writing of this report. This manuscript reflects the views of the authors and may not reflect the opinions or views of the NIH or ABCD consortium investigators. The ABCD data repository grows and changes over time.

The ABCD data used in this report came from https://nda.nih.gov/study.html?id=2313.

## References

1. Karcher NR. Psychotic-like experiences in childhood and early adolescence: Clarifying the construct and future directions. Schizophr Res [Internet]. 2022 Aug 1 [cited 2023 Aug 10];246:205–6. Available from: https://pubmed.ncbi.nlm.nih.gov/35809352/

2. Staines L, Healy C, Coughlan H, Clarke M, Kelleher I, Cotter D, et al. Psychotic experiences in the general population, a review; definition, risk factors, outcomes and interventions. Psychol Med [Internet]. 2022 Nov 25 [cited 2024 Feb 29];52(15):3297. Available from: /pmc/articles/PMC9772919/

3. Fisher HL, Caspi A, Poulton R, Meier MH, Houts R, Harrington H, et al. Specificity of childhood psychotic symptoms for predicting schizophrenia by 38 years of age: a birth cohort study. Psychol Med. 2013/01/11. 2013;43(10):2077–86.

4. Poulton R, Caspi A, Moffitt TE, Cannon M, Murray R, Harrington H. Children’s self-reported psychotic symptoms and adult schizophreniform disorder: a 15-year longitudinal study. Arch Gen Psychiatry [Internet]. 2000;57(11):1053–8. Available from: https://www.ncbi.nlm.nih.gov/pubmed/11074871

5. Poulton R, Caspi A, Moffitt TE, Cannon M, Murray R, Harrington H. Children’s Self-Reported Psychotic Symptoms and Adult Schizophreniform Disorder: A 15-Year Longitudinal Study. Arch Gen Psychiatry [Internet]. 2000 Nov 1 [cited 2023 Jun 4];57(11):1053–8. Available from: https://jamanetwork.com/journals/jamapsychiatry/fullarticle/481671

6. Schoorl J, Barbu M, Shen X, … MHT, 2021 undefined. Grey and white matter associations of psychotic-like experiences in a general population sample (UK Biobank). nature.com [Internet]. [cited 2023 Apr 17]; Available from: https://www.nature.com/articles/s41398-020-01131-7

7. Evermann U, Gaser C, Besteher B, Langbein K, Nenadić I. Cortical Gyrification, Psychotic-Like Experiences, and Cognitive Performance in Nonclinical Subjects. Schizophr Bull [Internet]. 2020 Dec 1 [cited 2023 Apr 17];46(6):1524–34. Available from: https://academic.oup.com/schizophreniabulletin/article/46/6/1524/5874167

8. Satterthwaite TD, Wolf DH, Calkins ME, Vandekar SN, Erus G, Ruparel K, et al. Structural Brain Abnormalities in Youth With Psychosis Spectrum Symptoms. JAMA Psychiatry. 2016/03/17. 2016;73(5):515–24.

9. Karcher NR, Barch DM, Avenevoli S, Savill M, Huber RS, Simon TJ, et al. Assessment of the Prodromal Questionnaire-Brief Child Version for Measurement of Self-reported Psychoticlike Experiences in Childhood. JAMA Psychiatry. 2018/06/07. 2018;75(8):853–61.

10. Karcher NR, Loewy RL, Savill M, Avenevoli S, Huber RS, Makowski C, et al. Persistent and distressing psychotic-like experiences using adolescent brain cognitive development^SM^ study data. Mol Psychiatry. 2021;1–12.

11. Romer AL, Pizzagalli DA. Associations between Brain Structural Alterations, Executive Dysfunction, and General Psychopathology in a Healthy and Cross-Diagnostic Adult Patient Sample. Biological Psychiatry Global Open Science [Internet]. 2021; Available from: https://www.sciencedirect.com/science/article/pii/S2667174321000483

12. Karcher NR, Merchant J, Pine J, Kilciksiz CM. Cognitive Dysfunction as a Risk Factor for Psychosis. Curr Top Behav Neurosci [Internet]. 2023 [cited 2024 Feb 29];63:173–203. Available from: https://link.springer.com/chapter/10.1007/7854_2022_387

13. Gur RC, Calkins ME, Satterthwaite TD, Ruparel K, Bilker WB, Moore TM, et al. Neurocognitive growth charting in psychosis spectrum youths. JAMA Psychiatry. 2014;71(4):366–74.

14. Dickson H, Cullen AE, Jones R, Reichenberg A, Roberts RE, Hodgins S, et al. Trajectories of cognitive development during adolescence among youth at-risk for schizophrenia. J Child Psychol Psychiatry. 2018/04/24. 2018;59(11):1215–24.

15. Tamnes CK, Herting MM, Goddings AL, Meuwese R, Blakemore SJ, Dahl RE, et al. Development of the Cerebral Cortex across Adolescence: A Multisample Study of Inter-Related Longitudinal Changes in Cortical Volume, Surface Area, and Thickness. Journal of Neuroscience [Internet]. 2017 Mar 22 [cited 2024 Mar 27];37(12):3402–12. Available from: https://www.jneurosci.org/content/37/12/3402

16. Thorup AAE, Hemager N, Bliksted VF, Greve AN, Ohland J, Wilms M, et al. The Danish High-Risk and Resilience Study— VIA 15 – A Study Protocol for the Third Clinical Assessment of a Cohort of 522 Children Born to Parents Diagnosed With Schizophrenia or Bipolar Disorder and Population-Based Controls. Front Psychiatry [Internet]. 2022 Apr 4 [cited 2024 Mar 28];13:809807. Available from: www.frontiersin.org

17. Collins MA, Ji JL, Chung Y, Lympus CA, Afriyie-Agyemang Y, Addington JM, et al. Accelerated cortical thinning precedes and predicts conversion to psychosis: The NAPLS3 longitudinal study of youth at clinical high-risk. Mol Psychiatry [Internet]. 2023 Mar 1 [cited 2024 Mar 29];28(3):1182. Available from: /pmc/articles/PMC10005940/

18. Cannon TD, Chung Y, He G, Sun D, Jacobson A, van Erp TG, et al. Progressive reduction in cortical thickness as psychosis develops: a multisite longitudinal neuroimaging study of youth at elevated clinical risk. Biol Psychiatry. 2014/07/19. 2015;77(2):147–57.

19. Pantelis C, Velakoulis D, McGorry PD, Wood SJ, Suckling J, Phillips LJ, et al. Neuroanatomical abnormalities before and after onset of psychosis: a cross-sectional and longitudinal MRI comparison. Lancet [Internet]. 2003 Jan 25 [cited 2023 Apr 1];361(9354):281–8. Available from: https://pubmed.ncbi.nlm.nih.gov/12559861/

20. Merritt K, Luque Laguna P, Irfan A, David AS. Longitudinal Structural MRI Findings in Individuals at Genetic and Clinical High Risk for Psychosis: A Systematic Review. Front Psychiatry. 2021 Feb 2;12:49.

21. van Os J, Linscott RJ, Myin-Germeys I, Delespaul P, Krabbendam L. A systematic review and meta-analysis of the psychosis continuum: evidence for a psychosis proneness-persistence-impairment model of psychotic disorder. Psychol Med [Internet]. 2009;39(2):179–95. Available from: https://www.ncbi.nlm.nih.gov/pubmed/18606047

22. Karcher NR, Paul SE, Johnson EC, Hatoum AS, Baranger DAA, Agrawal A, et al. Psychotic-like Experiences and Polygenic Liability in the Adolescent Brain Cognitive Development Study. Biol Psychiatry Cogn Neurosci Neuroimaging. 2022 Jan 1;7(1):45–55.

23. Karcher NR, Schiffman J, Barch DM. Environmental Risk Factors and Psychotic-like Experiences in Children Aged 9–10. J Am Acad Child Adolesc Psychiatry [Internet]. 2021;60(4):490–500. Available from: https://www.sciencedirect.com/science/article/pii/S0890856720303920

24. Glenthøj LB, Hjorthøj C, Kristensen TD, Davidson CA, Nordentoft M. The effect of cognitive remediation in individuals at ultra-high risk for psychosis: a systematic review. NPJ Schizophr [Internet]. 2017 Dec 1 [cited 2024 Aug 5];3(1):20. Available from: /pmc/articles/PMC5441569/

25. Karcher NR, Modi H, Kochunov P, Gao S, Barch DM. Regional Vulnerability Indices in Youth With Persistent and Distressing Psychoticlike Experiences. JAMA Netw Open [Internet]. 2023 Nov 1 [cited 2024 Mar 27];6(11):e2343081–e2343081. Available from: https://jamanetwork.com/journals/jamanetworkopen/fullarticle/2811762

26. Chapman LJ, Chapman JP, Kwapil TR, Eckblad M, Zinser MC. Putatively psychosis-prone subjects 10 years later. J Abnorm Psychol [Internet]. 1994;103(2):171–83. Available from: https://www.ncbi.nlm.nih.gov/pubmed/8040487

27. Weintraub S, Dikmen SS, Heaton RK, Tulsky DS, Zelazo PD, Bauer PJ, et al. Cognition assessment using the NIH Toolbox. Neurology [Internet]. 2013;80(11 Suppl 3):S54–64. Available from: https://www.ncbi.nlm.nih.gov/pubmed/23479546

28. Fischl B, Sereno MI, Dale AM. Cortical surface-based analysis. II: Inflation, flattening, and a surface-based coordinate system. Neuroimage. 1999/02/05. 1999;9(2):195–207.

29. Chen CH, Gutierrez ED, Thompson W, Panizzon MS, Jernigan TL, Eyler LT, et al. Hierarchical genetic organization of human cortical surface area. Science (1979). 2012/03/31. 2012;335(6076):1634–6.

30. Fischl B, Dale AM. Measuring the thickness of the human cerebral cortex from magnetic resonance images. Proc Natl Acad Sci U S A. 2000/09/14. 2000;97(20):11050–5.

31. Hagler Jr DJ, Hatton S, Cornejo MD, Makowski C, Fair DA, Dick AS, et al. Image processing and analysis methods for the Adolescent Brain Cognitive Development Study. Neuroimage. 2019;202:116091.

32. Beer JC, Tustison NJ, Cook PA, Davatzikos C, Sheline YI, Shinohara RT, et al. Longitudinal ComBat: A method for harmonizing longitudinal multi-scanner imaging data. Neuroimage. 2020 Oct 15;220:117129.

33. Ripke S, Walters JTR, O’Donovan MC. Mapping genomic loci prioritises genes and implicates synaptic biology in schizophrenia. medRxiv [Internet]. 2020;2020.09.12.20192922. Available from: https://www.medrxiv.org/content/medrxiv/early/2020/09/13/2020.09.12.20192922.full.pdf

34. Bigdeli TB, Genovese G, Georgakopoulos P, Meyers JL, Peterson RE, Iyegbe CO, et al. Contributions of common genetic variants to risk of schizophrenia among individuals of African and Latino ancestry. Mol Psychiatry [Internet]. 2020;25(10):2455–67. Available from: 10.1038/s41380-019-0517-y

35. Ge T, Chen C, Ni Y, Feng Y, communications JSN, 2019 undefined. Polygenic prediction via Bayesian regression and continuous shrinkage priors. nature.com [Internet]. [cited 2023 Jun 23]; Available from: https://www.nature.com/articles/s41467-019-09718-5

36. Martin AR, Kanai M, Kamatani Y, Okada Y, Neale BM, Daly MJ. Clinical use of current polygenic risk scores may exacerbate health disparities. Nat Genet [Internet]. 2019;51(4):584–91. Available from: http://europepmc.org/abstract/MED/30926966

37. Karcher NR, Merchant J, Pine J, Kilciksiz CM. Cognitive Dysfunction as a Risk Factor for Psychosis. Curr Top Behav Neurosci [Internet]. 2023 [cited 2023 Apr 10];63:173–203. Available from: https://link.springer.com/chapter/10.1007/7854_2022_387

38. Fusar-Poli P, Tantardini M, De Simone S, Ramella-Cravaro V, Oliver D, Kingdon J, et al. Deconstructing vulnerability for psychosis: Meta-analysis of environmental risk factors for psychosis in subjects at ultra high-risk. European Psychiatry. 2017 Feb 1;40:65–75.

39. Linscott RJ, van Os J. An updated and conservative systematic review and meta-analysis of epidemiological evidence on psychotic experiences in children and adults: on the pathway from proneness to persistence to dimensional expression across mental disorders. Psychol Med [Internet]. 2013;43(6):1133–49. Available from: https://www.ncbi.nlm.nih.gov/pubmed/22850401

40. Martin AR, Kanai M, Kamatani Y, Okada Y, Neale BM, Daly MJ. Clinical use of current polygenic risk scores may exacerbate health disparities. Nat Genet. 2019 Apr 1;51(4):584–91.

