## Supplemental Methods and Results for "Longitudinal Trajectories of Cognition and Neural Metrics as Predictors of Persistent Distressing Psychotic-Like Experiences Across Middle Childhood and Early Adolescence"

#### Supplemental Materials

##### Table of Contents

Supplemental Methods…..…………………………………………………...……….. Pages 2-5
Participants……………….……………………..…………………..……...…..…...Page 2
Measures……………………………………..……………………………...…..Pages 2-5
Statistical Analyses………….…………….………………….……..…..……...Page 5
Supplemental Results……………….………………….………………..……...…..Page 5-6
Supplemental References……………………….………………..……...…..…...Pages 7-8

Supplemental Figures……………………..………..…………………………Pages 9-14

#### Supplemental Methods

##### Participants

The ABCD study aimed to recruit a sample reflecting the demographic variation of the U.S. population, recruiting children using probability sampling from both public and private elementary schools. Study-wide exclusionary criteria were as follows: child not fluent in English, MRI contraindication (e.g., irremovable ferromagnetic implants or dental appliances, claustrophobia, pregnant), major neurological disorder, gestational age less than 28 weeks or birthweight less than 1,200 grams, history of traumatic brain injury, or had a current diagnosis of schizophrenia, autism spectrum disorder (moderate, severe), mental retardation/intellectual disability, or alcohol/substance use disorder.[1–3] Parents provided written informed consent and all children provided assent.

##### Measures

***Pubertal Status.*** Pubertal status was assessed via the caregiver-reported Pubertal Development Scale, with the current study examining the mean of five questions regarding changes in physical characteristics (e.g., height, body hair, skin), with each question rated on a 4-point scale from 1=has not begun yet to 4=seems complete.[4]

***Youth-Reported Internalizing and Externalizing Symptoms.*** Youth-reported internalizing and externalizing psychopathology was assessed at the 6-month follow-up (and every six months after including the 4-year follow-up) using an abbreviated form of the Youth Self Report, the child-rated Brief Problem Monitor for Youth (BPM-Y). Analyses examined raw internalizing and externalizing scales.[27]

***MRI Analyses***. Structural neuroimaging processing was completed using FreeSurfer version 5.3.0 through standardized processing pipelines.[5] Participants that did not pass FreeSurfer Quality Control measure (i.e., at least one T1 scan that passed all quality control metrics) were excluded from analyses. Cortical reconstruction and volumetric segmentation was performed by the ABCD Data Acquisition and Integration Core using the FreeSurfer image analysis suite (http://surfer.nmr.mgh.harvard.edu/). This pre-processing includes removal of non-brain tissue using a hybrid watershed/surface deformation procedure,[6] automated Talairach transformation, segmentation of the subcortical white matter and deep gray matter volumetric structures, intensity normalization, tessellation of the gray/white matter boundary, automated topology correction, and surface deformation following intensity gradients. Images were registered to an atlas, which was based on individual cortical folding patterns to match cortical geometry across subjects. The following global and regional structural MRI metrics were examined: *global:* intracranial, total cortical, and total subcortical volume; total surface area, and total cortical thickness.

**Environmental Risk Factors.** A number of environmental risk factors were retrieved based on previous research and available data at baseline.[7] Perception of neighborhood safety was calculated as a summation of three parent-rated questions (i.e., “I feel safe walking in my neighborhood, day or night”; “Violence is not a problem in my neighborhood”; “My neighborhood is safe from crime”; each was rated on a scale from 1-5, 1=strongly disagree, 5=strongly agree). Next, total drug offenses crime exposure information was obtained from the Uniform Crime Report from FBI, compiled by Inter-university Consortium for Political and Social Research,[8] averaged from 2010-2012 to create stable county-level estimates. Overall deprivation was defined as the Area Deprivation Index (ADI) national percentile scores for the current residential address at baseline, calculated from the 2011-2015 American Community Survey 5-year summary.[9] We also examined the proportion of individuals living in poverty (-125% of poverty level) and number of years at current residence. Estimates of lead exposure risk were obtained by first geocoding the participant’s address at the census tract-level and then calculating risk scores based on data obtained from vox.com (https://www.vox.com/a/lead-exposure-risk-map). Estimated lead exposure risk scores (1-10, 10 being the most at risk) were calculated using proportion of individuals living in poverty and average age of the home (see *Deprivation* section above).[7]

**Genotyping, Quality Control, and Imputation.** The ABCD sample was genotyped using Rutgers University Cell and DNA repository, who genotyped saliva samples on the Smokescreen array.[10] Genotyped calls were aligned to GRCh37 (hg19), and all individuals self-reporting ancestral origins (i.e., self-reported race) other than European were excluded due to our necessary use of summary statistics based on GWAS conducted in European ancestral samples and evidence that the predictive utility of polygenic risk scores suffers when applied across ancestral origins (although exploratory analyses examined an schizophrenia PGS, created using summary statistics from a schizophrenia GWAS with individuals of African ancestry, in a subsample of individuals of African descent).[11]

The following preprocessing steps were conducted with the Ricopili pipeline.[12] In short, SNPs with call rates ≥ 0·95 and MAF ≥ 1% were retained. Individuals with high rates of missingness (>5%) and autosomal heterozygosity deviation ($F_{HET}$) outside of ± 2 SD were removed. After sample quality control (QC), SNPs were further filtered to call rate ≥ 0·98 and Hardy-Weinberg p-values > $10^{-6}$ (founders only), which yielded 372,342 SNPs. In order to reconcile mismatches, sex checks were conducted with follow-up.

Individuals whose data passed the first phase of QC were then checked for relatedness--both known and cryptic--and Mendelian errors were resolved. Next, using data from unrelated individuals (pi-hat ≤ 0.2) and an LD pruned set of common (MAF>0.05) and non-palindromic SNPs (and excluding MHC and chromosome eight inversion region), principal components analysis (PCA) was performed in EIGENSTRAT using the cosmopolitan 1000 Genomes Project phase 3 data. Only those individuals whose data aligned with ancestral non-Hispanic European ancestry were retained, yielding a sample of 5,556. After selection, a final ancestrally-informative PCA was conducted, and the first ten PCs were projected from founders to other relatives. Imputation to Haplotype Reference Consortium (HRC) data for Europeans (and CAPPA for African Americans, see Supplemental Results below) was conducted using strictly QCed SNPs on the Michigan Imputation Server, yielding 39,127,678 SNPs. Dosage data were converted to hard-call genotypes using Plink, and only SNPs with imputation $r^2$ scores ≥ 0.3 were used to create polygenic scores. For the main analyses of the manuscript, the final sample of individuals passing all QC metrics was n=4,650.

**Polygenic Scores.** Polygenic scores (PGS) were generated using the Polygenic Risk Score-Continuous Shrinkage (PRS-CS) software package.[13] We used the PRS-CS "auto" function, which employs a fully Bayesian approach such that the global shrinkage parameter, $\phi$, is automatically learned from data. This method does not prune SNPs based on p-value threshold or perform clumping for independent SNPs; instead, it assumes a general distribution of effect sizes across the genome and accounts for linkage disequilibrium (LD) between SNPs using an external LD reference panel (1000 Genomes Phase 3 European samples). Our models were trained using summary statistics from the Schizophrenia GWAS (Psychiatric Genetics Consortium, N=69,369 cases + 236,642 controls).[14]

#### Statistical Analysis

We followed up the a priori PLE groups by examining latent profile analyses (LPA; using tidyLPA and mclust packages).[18,19] Latent profile models with one through five profiles were estimated using PLE distress scores at the five time points. Models were evaluated based on model fit indices, and when there was disagreement between fit indices in terms of which model showed the best fit, model selection was based on the lowest BIC [20]. Latent growth curves compared extracted latent classes on trajectories of cognitive and neural data. We also re-run analyses presented in the manuscript including imputed data (Supplemental Table 11).

### Supplemental Results

#### Latent Class Analyses

Latent profile analyses revealed two classes showing stable PLE trends over time: low and high profiles (Supplemental Figure 5; Supplemental Table 8). Consistent with the a priori PLE group results, the high PLE latent profile was associated with worsening cognitive and impaired structural neural metric trajectories over time across all metrics (Supplemental Table 9).

#### Supplemental References


1. Garavan H, Bartsch H, Conway K, … ADD cognitive, 2018 undefined. Recruiting the ABCD sample: Design considerations and procedures. *Elsevier*. Accessed June 8, 2023. https://www.sciencedirect.com/science/article/pii/S1878929317301809
2. Karcher NR, Barch DM. The ABCD study: understanding the development of risk for mental and physical health outcomes. *Neuropsychopharmacology 2020 46:1*. 2020;46(1):131-142. doi:10.1038/s41386-020-0736-6
3. Barch DM, Albaugh MD, Baskin-Sommers A, et al. Demographic and mental health assessments in the adolescent brain and cognitive development study: Updates and age-related trajectories. *Dev Cogn Neurosci*. 2021;52:101031. doi:10.1016/J.DCN.2021.101031
4. Petersen AC, Crockett L, Richards M, Boxer A. A self-report measure of pubertal status: Reliability, validity, and initial norms. *J Youth Adolesc*. 1988;17(2):117-133.
5. Hagler Jr DJ, Hatton S, Cornejo MD, et al. Image processing and analysis methods for the Adolescent Brain Cognitive Development Study. *Neuroimage*. 2019;202:116091.
6. Segonne F, Dale AM, Busa E, et al. A hybrid approach to the skull stripping problem in MRI. *Neuroimage*. 2004;22(3):1060-1075. doi:10.1016/j.neuroimage.2004.03.032
7. Karcher NR, Schiffman J, Barch DM. Environmental Risk Factors and Psychotic-like Experiences in Children Aged 9–10. *J Am Acad Child Adolesc Psychiatry*. 2021;60(4):490-500. doi:https://doi.org/10.1016/j.jaac.2020.07.003
8. Investigation USD of JusticeO of JProgramsFB of. Uniform Crime Reporting Program Data: County-Level Detailed Arrest and Offense Data, United States, 2010. Published online 2014. doi:10.3886/ICPSR33523.v2
9. Kind AJH, Jencks S, Brock J, et al. Neighborhood socioeconomic disadvantage and 30-day rehospitalization: a retrospective cohort study. *Ann Intern Med*. 2014;161(11):765-774.
10. Baurley JW, Edlund CK, Pardamean CI, Conti D V, Bergen AW. Smokescreen: a targeted genotyping array for addiction research. *BMC Genomics*. 2016;17(1):145.
11. Bogdan R, Baranger DAA, Agrawal A. Polygenic Risk Scores in Clinical Psychology: Bridging Genomic Risk to Individual Differences. *Annu Rev Clin Psychol*. 2018;14:119-157. doi:10.1146/annurev-clinpsy-050817-084847
12. Lam M, Awasthi S, Watson HJ, et al. RICOPILI: Rapid Imputation for COnsortias PIpeLIne. *Bioinformatics*. 2020;36(3):930-933.
13. Ge T, Chen C, Ni Y, Feng Y, communications JSN, 2019 undefined. Polygenic prediction via Bayesian regression and continuous shrinkage priors. *nature.com*. Accessed June 23, 2023. https://www.nature.com/articles/s41467-019-09718-5
14. Consortium SWG of the PG. Biological insights from 108 schizophrenia-associated genetic loci. *Nature*. 2014;511(7510):421-427. doi:10.1038/nature13595
15. Lee PH, Anttila V, Won H, et al. Genomic relationships, novel loci, and pleiotropic mechanisms across eight psychiatric disorders. *Cell*. 2019;179:1469-1482. doi:10.1016/j.cell.2019.11.020
16. Legge SE, Jones HJ, Kendall KM, et al. Association of Genetic Liability to Psychotic Experiences With Neuropsychotic Disorders and Traits. *JAMA Psychiatry*. 2019;76(12):1256-1265. doi:10.1001/jamapsychiatry.2019.2508

17. Lee JJ, Wedow R, Okbay A, et al. Gene discovery and polygenic prediction from a genome-wide association study of educational attainment in 1.1 million individuals. *Nat Genet*. 2018;50(8):1112-1121.
18. Scrucca L, Fop M, Murphy T, journal ARTR, 2016 undefined. mclust 5: clustering, classification and density estimation using Gaussian finite mixture models. *ncbi.nlm.nih.govL Scrucca, M Fop, TB Murphy, AE RafteryThe R journal, 2016•ncbi.nlm.nih.gov*. Accessed August 28, 2024. https://www.ncbi.nlm.nih.gov/pmc/articles/PMC5096736/
19. Rosenberg JM, Beymer PN, Anderson DJ, Lissa C j. van, Schmidt JA. tidyLPA: An R Package to Easily Carry Out Latent Profile Analysis (LPA) Using Open-Source or Commercial Software. *J Open Source Softw*. 2019;3(30):978. doi:10.21105/JOSS.00978
20. Chakrabarti A, Ghosh JK. AIC, BIC and Recent Advances in Model Selection. *Philosophy of Statistics*. Published online January 1, 2011:583-605. doi:10.1016/B978-0-444-51862-0.50018-6

**Supplemental Figure Captions**

**Supplemental Figure 1.** Plots illustrating individual trajectories for each of the PLEs groups across the three time points for cognitive metrics.
**Supplemental Figure 2.** Plots illustrating individual trajectories for each of the PLEs groups across the three time points for neural metrics.
**Supplemental Figure 3.** Plots illustrating distributions of cognitive metric variables for each of the PLEs groups at each time point.
**Supplemental Figure 4.** Plots illustrating distributions of neural metric variables for each of the PLEs groups at each time point.
**Supplemental Figure 5.** Plots illustrate mean levels of psychotic-like experiences (PLE) for each of the two PLE latent profiles.

*Supplemental Figure 1.*

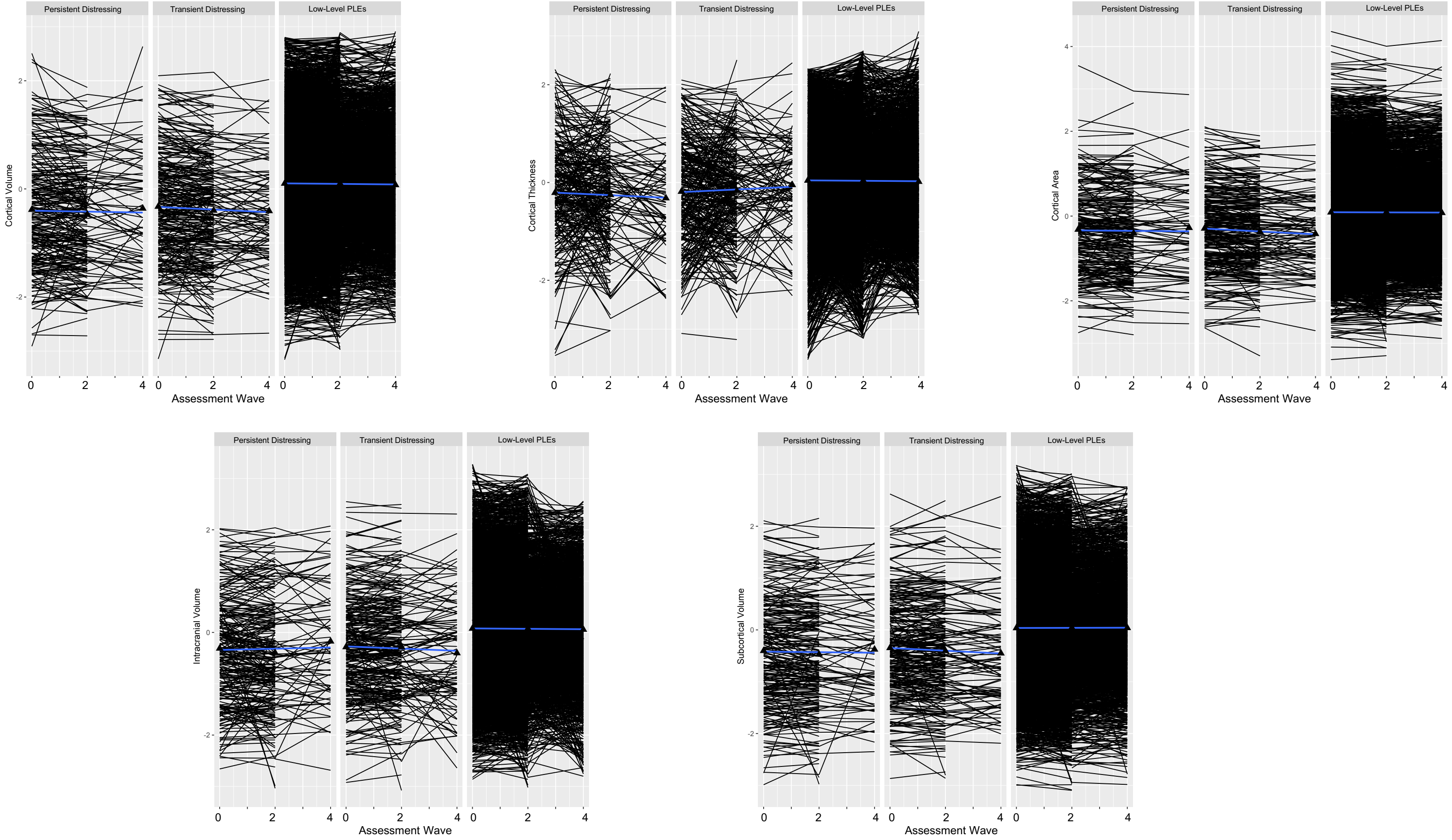

*Supplemental Figure 2.*

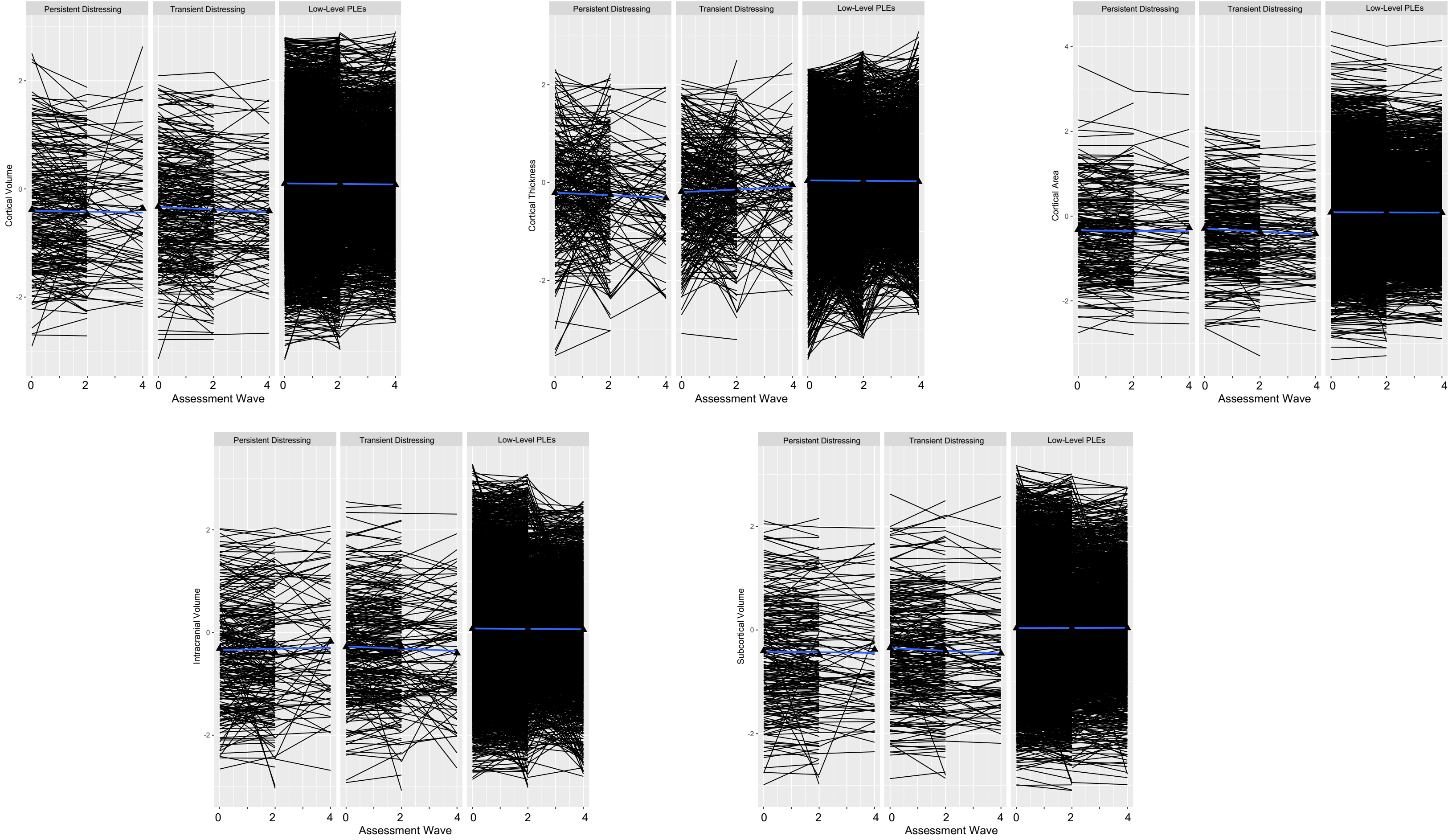

*Supplemental Figure 3.*

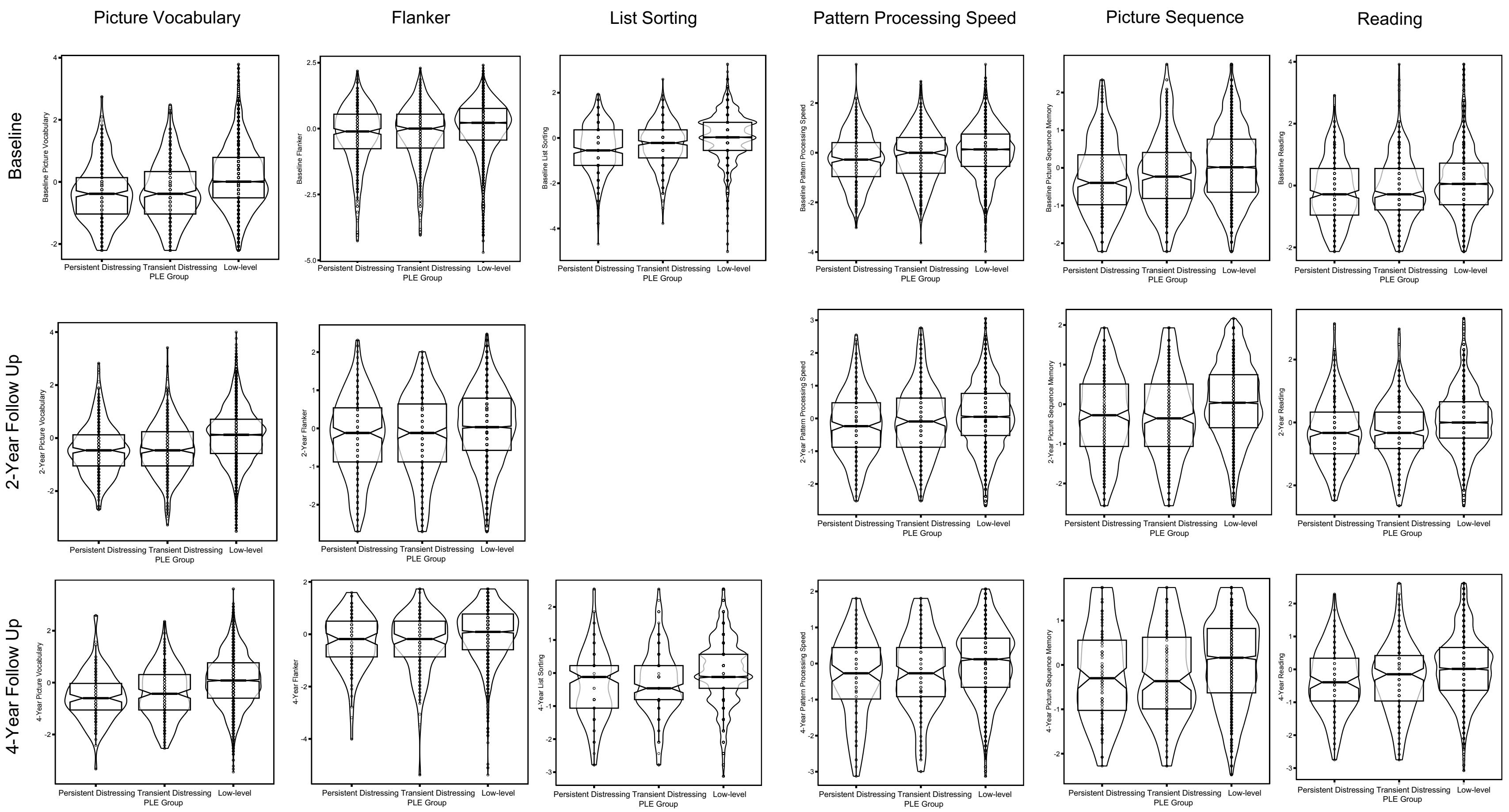

*Supplemental Figure 4.*

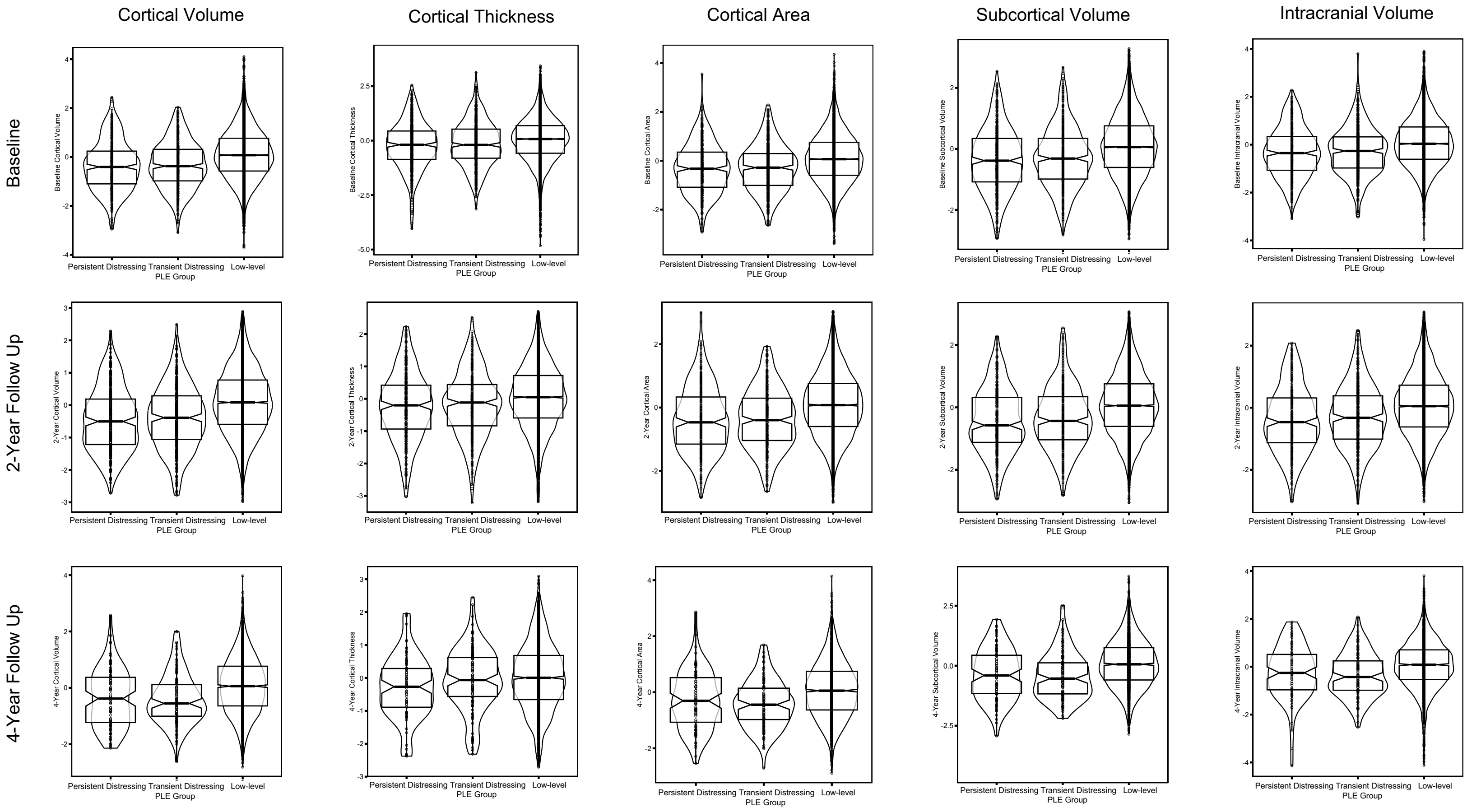

*Supplemental Figure 5.*

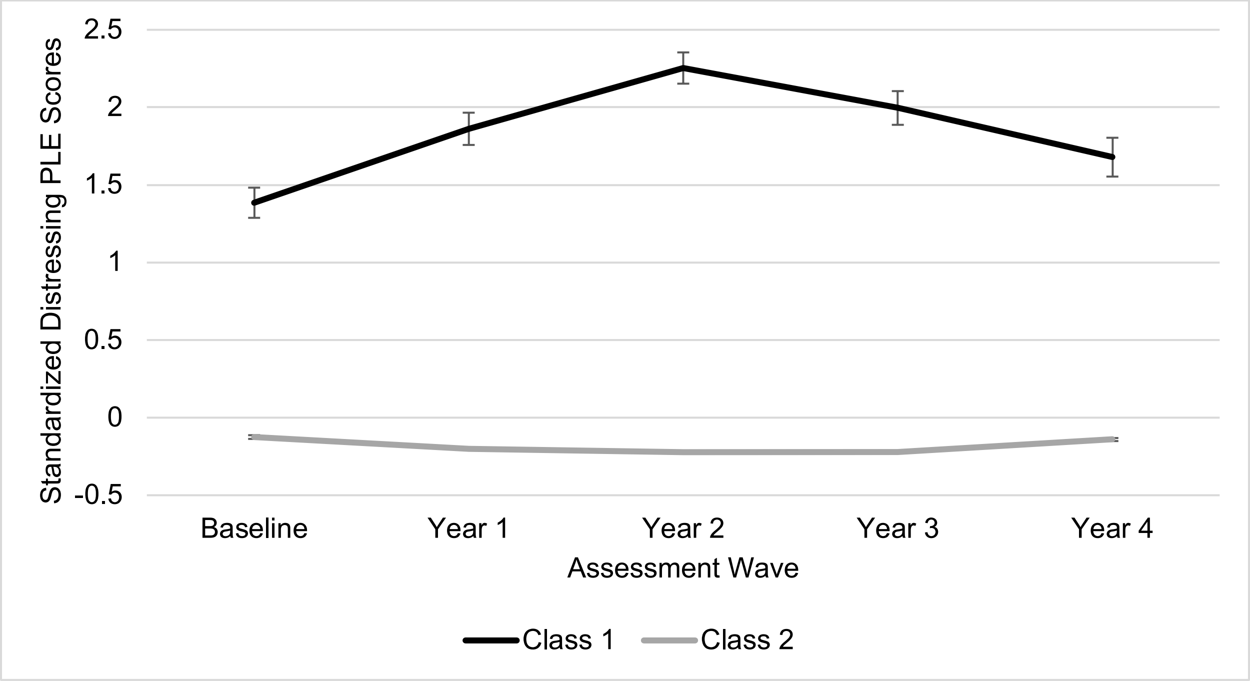
